# Identifying SARS-CoV-2 Antiviral Compounds by Screening for Small Molecule Inhibitors of Nsp12/7/8 RNA-dependent RNA Polymerase

**DOI:** 10.1101/2021.04.07.438807

**Authors:** Agustina P. Bertolin, Florian Weissmann, Jingkun Zeng, Viktor Posse, Jennifer C. Milligan, Berta Canal, Rachel Ulferts, Mary Wu, Lucy S. Drury, Michael Howell, Rupert Beale, John F.X. Diffley

## Abstract

The coronavirus disease 2019 (COVID-19) global pandemic has turned into the largest public health and economic crisis in recent history impacting virtually all sectors of society. There is a need for effective therapeutics to battle the ongoing pandemic. Repurposing existing drugs with known pharmacological safety profiles is a fast and cost-effective approach to identify novel treatments. The COVID-19 etiologic agent is the severe acute respiratory syndrome coronavirus 2 (SARS-CoV-2), a single-stranded positive-sense RNA virus. Coronaviruses rely on the enzymatic activity of the replication-transcription complex (RTC) to multiply inside host cells. The RTC core catalytic component is the RNA-dependent RNA polymerase (RdRp) holoenzyme. The RdRp is one of the key druggable targets for CoVs due to its essential role in viral replication, high degree of sequence and structural conservation and the lack of homologs in human cells. Here, we have expressed, purified and biochemically characterised active SARS-CoV-2 RdRp complexes. We developed a novel fluorescence resonance energy transfer (FRET)-based strand displacement assay for monitoring SARS-CoV-2 RdRp activity suitable for a high-throughput format. As part of a larger research project to identify inhibitors for all the enzymatic activities encoded by SARS-CoV-2, we used this assay to screen a custom chemical library of over 5000 approved and investigational compounds for novel SARS-CoV-2 RdRp inhibitors. We identified 3 novel compounds (GSK-650394, C646 and BH3I-1) and confirmed suramin and suramin-like compounds as *in vitro* SARS-CoV-2 RdRp activity inhibitors. We also characterised the antiviral efficacy of these drugs in cell-based assays that we developed to monitor SARS-CoV-2 growth.

## Introduction

The COVID-19 outbreak caused by SARS-CoV-2 is considered to be the most severe global public health crisis since the influenza pandemic in 1918 (1, 2). As of February 2021, there were more than 100 million confirmed cases and 2.5 million deaths globally (3). Remdesivir is currently the only antiviral drug approved for COVID-19 treatment. Remdesivir is a nucleoside analog that acts as a small molecule inhibitor of an essential coronavirus enzyme complex, the RNA-dependent RNA polymerase (4). Nucleoside analogues are one of the largest and most effective classes of small molecule drugs against viruses like human immunodeficiency virus, hepatitis B virus and herpesviruses (5, 6). The efficacy of remdesivir as a COVID-19 treatment remains controversial. While one clinical trial found remdesivir to shorten recovery time in COVID-19 patients, other clinical trials found no effect of remdesivir on recovery time or mortality rate (7–10). Besides remdesivir, several other clinically approved nucleoside analog compounds are currently under study for treating COVID-19. Molnupiravir (11, 12) and AT-527 (13) are the most promising candidates so far, while ritonavir (14), ribavirin (15), favipiravir (16) and sofosbuvir (6, 17) have not demonstrated significant antiviral effect against SARS-CoV-2 in laboratory or clinical settings.

Coronaviruses (CoVs) are enveloped single-stranded positive-sense RNA viruses that belong to the order *Nidovirales* (18). CoV replication and transcription occur in the cytoplasm of infected cells and are mediated by the replication-transcription complex (RTC) (18, 19). The core RTC is composed of i) the RNA-dependent RNA polymerase (RdRp) - a trimeric complex of the catalytic subunit nonstructural protein (nsp) 12 and two accessory factors nsp7 and nsp8 (20, 21)-; ii) the nsp13 helicase (22–24) and iii) several RNA-processing enzymes such as nsp14, a bifunctional enzyme with 3’-to-5’ exoribonuclease (ExoN) and N7-methyltransferase activities (25–27). Replication of coronaviruses is more complex than that of other RNA viruses due to their unusually large genomes (28). Replication of these large genomes requires enhanced RdRp processivity, which is promoted by the two nsp12 accessory subunits, nsp7 and nsp8 (20, 21, 28). Replication of large genomes also involves enzymatic functions that could decrease the high error rate typical of viral RNA polymerases to avoid detrimental fitness effects (29). Coronaviruses encode a 3’-to-5’ ExoN (nsp14) that can remove mis-incorporated nucleotides (30–35). This potential replication proofreading function may diminish the effectiveness of immediate chain terminator nucleoside analogues, since they can be removed by the nsp14 exonuclease when added to the nascent RNA chain (31, 36). On the other hand, remdesivir works as a delayed chain terminator: its addition to the growing RNA molecule results in termination only after incorporation of up to three additional nucleotides and hence shows some resistance to excision by the nsp14 exonuclease (37, 38).

RdRp has three characteristics that when combined position it as a key drug target for CoVs: i) it is essential for viral replication; ii) it has a high level of structural conservation among coronaviruses; and iii) it lacks a counterpart in human cells (18). However, due to the proofreading activity of nsp14 exonuclease, the potential of nucleoside analogues as antiviral drugs for COVID-19 treatment may be limited. Instead, identifying RdRp inhibitors which are not nucleoside analogues from currently approved or experimental drugs may lead to earlier and faster clinical trials, as these drugs have known pharmacokinetic and pharmacodynamic profiles and drug regimens (39).

Here we describe the expression and purification of active SARS-CoV-2 RdRp holoenzyme (nsp12/nsp7/nsp8). We also developed a FRET-based strand displacement assay suitable for high throughput screening to identify potential RdRp inhibitors using a custom chemical library of over 5000 compounds. We demonstrate that several non-nucleoside analogues potently block RdRp activity *in vitro* and one of them, GSK-650394, potently inhibits SARS-CoV-2 infectivity in a cell-based model of viral infection.

## Results

### Protein expression and purification

Coronavirus RdRp constitutes the catalytic core of the RTC and is composed of nsp12 in complex with two copies of nsp8 and one copy of nsp7 (nsp12/nsp8_2_/nsp7) (40). In order to maximise the chances of generating active RdRp in sufficient amounts for high throughput screening, we followed two protein expression strategies. First, we chose a eukaryotic expression system and expressed proteins in baculovirus-infected insect cells (*Spodoptera frugiperda*, Sf; **Figure 1A**). Using a similar approach to previous work with SARS-CoV-1 RdRp, we expressed and purified nsp12/nsp7-nsp8 complex in which the C-terminus of nsp12 was tagged with 3xFlag and nsp7 and nsp8 were fused with a His_6_ tag linker (7H8, similar to (21)). Purification by 3xFlag affinity, heparin affinity, and size exclusion chromatography yielded a near-homogeneous protein preparation (**Figure 1A**, lane 3: nsp12-F/7H8). To be able to obtain better yields, we transferred the His_6_-tag to the catalytic subunit nsp12 and produced a complex containing a His_6_-3xFlag tag on nsp12 (Sf nsp12-HF) and a neutral 6 amino acid (Gly-Gly-Ser)_2_-linker between nsp7 and nsp8 (7L8). Purification of this complex by affinity to Ni-NTA agarose, ion exchange and size exclusion chromatography resulted in ∼100x higher protein yields (**Supplementary Table S1**), albeit with less stoichiometric nsp7-nsp8 fusion protein (**Figure 1A**, lane 4: nsp12-HF/7L8). We also tried co-expression of individual nsp7 and nsp8 subunits with tagged nsp12. Purification of this complex resulted in sub-stoichiometric amounts of the nsp7 and nsp8 subunits (**Figure 1A**, lane 5: nsp12-HF/7/8). We additionally purified the nsp7-His_6_-nsp8 fusion protein and nsp12-His_6_-3xFlag in isolation (**Figure 1A**, lane 1: 7H8, lane 2: nsp12-HF) obtaining very good yield for 7H8 (**Supplementary Table S1**). In a second approach, we expressed the three proteins individually in *E. coli* as N-terminal His-SUMO fusion proteins (**Figure 1B**). In this system, the affinity tag and SUMO fusion can be removed after affinity purification by a SUMO-specific protease (41), leaving behind the same N-terminus as would be generated by viral protease-mediated polyprotein cleavage in infected cells. We expressed nsp7, nsp8 and nsp12 using this system and purified the proteins by affinity to Ni-NTA agarose, fusion protein removal, ion exchange and size exclusion chromatography (**Figure 1B**).

**Figure 1.**
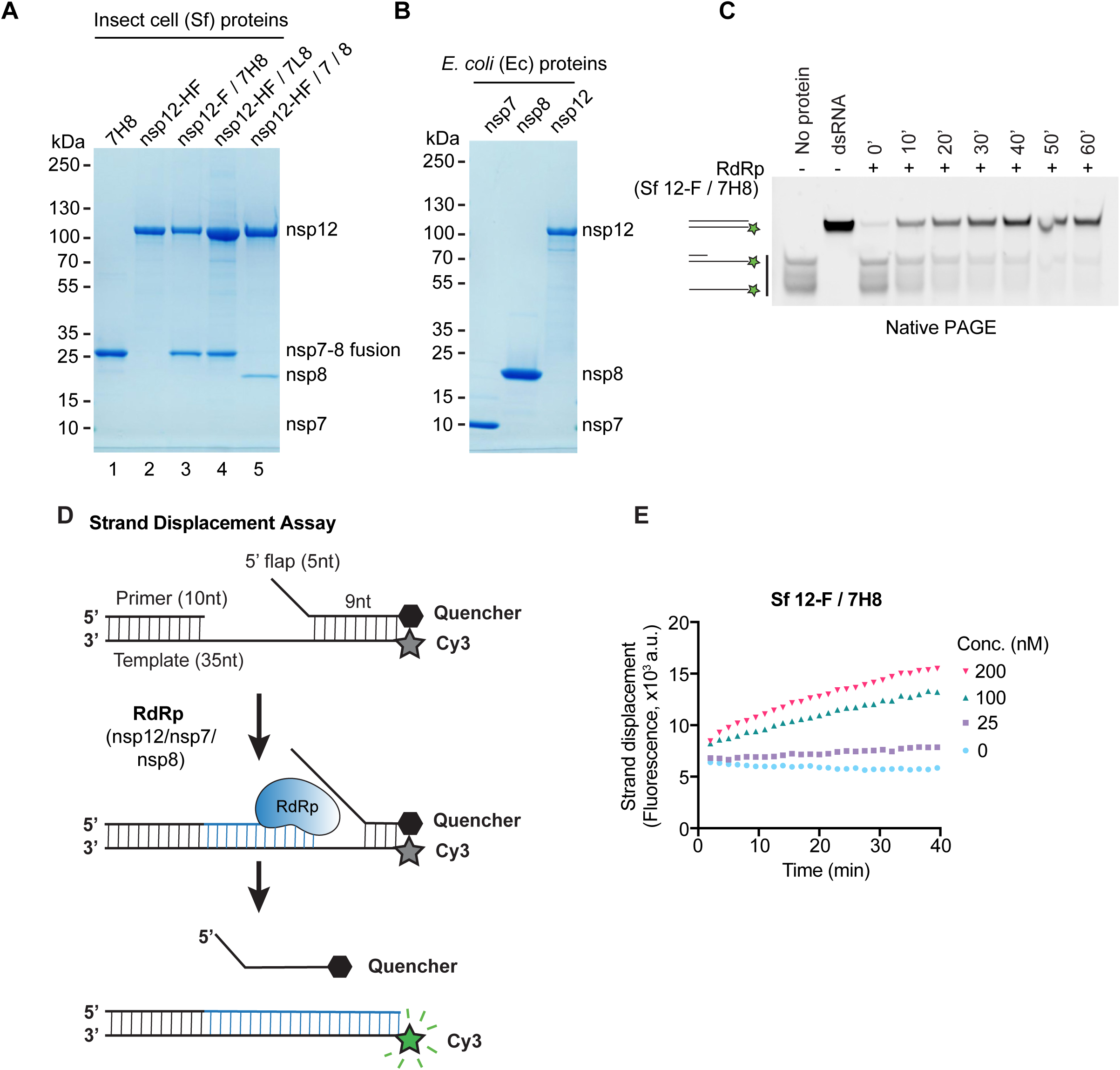
Development of a FRET-based SARS-CoV-2 RdRp strand displacement assay. (**A**) Purified SARS-CoV-2 RdRp proteins expressed in baculovirus-infected insect cells (*Spodoptera frugiperda*, Sf) analysed by SDS-PAGE and Coomassie staining. 7H8: nsp7-His_6_-nsp8, nsp12-HF: nsp12-His_6_-3xFlag, nsp12-F/7H8: nsp12-3xFlag/nsp7-His_6_-nsp8, nsp12-HF/7L8: nsp12-His_6_-3xFlag/nsp7-GGSGGS-nsp8, nsp12-HF/7/8: nsp12-His_6_-3xFlag/nsp7/nsp8. (**B**) Bacterially expressed and purified SARS-CoV-2 nsp7, nsp8 and nsp12 proteins analysed by SDS-PAGE and Coomassie staining. The proteins were expressed as 14His-SUMO fusion proteins in *E. coli*. 14His-SUMO was removed by a SUMO-specific protease during purification generating native N-termini. (**C**) Gel-based primer extension assay to test RNA-dependent RNA synthesis using the RdRp preparation Sf nsp12-F/7H8. The substrate consists of a 10 nt RNA primer annealed to the 3’ end of a 35 nt RNA template. The 5’ end of the template strand is labelled with a Cy3 fluorophore. Reaction products were analysed by native PAGE and visualisation of Cy3 fluorescence. Formation of duplex RNA by RdRp was observed over time. Controls: a preformed Cy3-labelled dsRNA with the same size as the reaction product (dsRNA), the primed substrate (No protein). (**D**) Schematic diagram illustrating the FRET-based RdRp strand displacement assay. The RNA substrate is composed of a Cy3 fluorophore-containing template strand, an annealed primer and an annealed quencher strand with a 5’ flap. RdRp activity synthesises RNA by extending the primer strand and displaces the quencher strand. The displaced quencher strand can no longer anneal to fully synthesised duplex RNA leading to an increase in Cy3 fluorescent signal. (**E**) FRET-based strand displacement assay using the indicated concentrations of Sf nsp12-F/7H8.

### Development of a FRET-based strand displacement assay for RdRp

To assess RNA-dependent RNA polymerisation activity we generated a primed RNA substrate by annealing a 35 nucleotide (nt) RNA template that contained a Cy3 fluorophore on its 5’ end with a 10 nt primer strand complementary to the 3’-end of the template. After incubation of RdRp with primed substrate and NTPs, reaction products were separated by native PAGE (**Figure 1C**). We tested in this assay the activity of the RdRp preparation from insect cells containing C-terminally tagged nsp12 and a co-expressed nsp7-nsp8 fusion protein with a His_6_-linker (Sf nsp12-F/7H8). This RdRp preparation was able to extend the primed RNA substrate to generate duplex RNA in a time-dependent manner, confirming RNA-dependent RNA polymerisation activity (**Figure 1C**).

We developed a variation of this primer-extension assay that can measure activity directly in solution, without requiring running products in a gel (**Supplementary Figure S1A,B**). As in Figure 1C, the Cy3-labelled primed RNA substrate is extended by the RdRp complex and generates duplex RNA. Then, after 60 minutes, we added a 14 nt RNA strand partially complementary to the template strand with a quencher molecule at its 3’ end. If RdRp has extended the primer, this quencher strand will *not* anneal to the template strand and will not be able to quench Cy3 fluorescence (**Supplementary Figure S1A**). We tested nsp12-F/7H8 in this assay and found that Cy3 fluorescence was greatly increased when RdRp was included in the reaction and the presence of Mn^2+^ enhanced RdRp activity compared to Mg^2+^ alone (**Supplementary Figure S1B**), which is in line with a published SARS-CoV-1 nsp12 enzymatic characterization (42).

None of the primer-extension assays described above (**Figure 1C** and **Supplementary Figure S1A and B**) are amenable to accurate high throughput screening (HTS) as they involve multiple steps and rely only on end point values. Therefore, we designed a FRET-based assay suitable for HTS based on RNA synthesis-coupled strand displacement activity (**Figure 1D**). Strand displacement refers to the ability of certain DNA/RNA polymerases to displace downstream non-template strands from the template strand while polymerising nucleotides (43, 44) (**Figure 1D**). The RNA substrate was constructed by annealing the primed 35 nt RNA template with the 14 nt quencher strand (**Figure 1D**). This structure places the Cy3 fluorophore in close proximity to the quencher localised on the opposite strand. As RdRp elongates the primer, it displaces the downstream quencher strand producing a fluorescent signal. As the final product is an RNA duplex, the quencher strand is prevented from reannealing (**Figure 1D**). When Sf nsp12-F/7H8 was incubated with the strand displacement substrate, fluorescence increased near-linearly with time and was dependent on enzyme concentration (**Figure 1E**). The presence of Mn^2+^ was not required but again greatly enhanced RdRp activity compared to Mg^2+^ alone (**Supplementary Figure S1C-E**). The fluorescence increase was dependent on NTPs (**Supplementary Figure S1F**) suggesting that i) there were no contaminating nucleases in the reactions that could also have resulted in freeing the Cy3 fluorophore from the quencher and ii) RdRp polymerisation was driving strand displacement of the quencher strand. We tested our different RdRp preparations from Figure 1A and 1B using the FRET-based strand displacement assay in a concentration range of 25 nM - 400 nM (**Figure 2**). The C-terminally tagged nsp12 co-expressed with nsp7-nsp8 fusion protein with His linker (Sf nsp12-F/7H8, **Figure 2A**) was slightly more active than Sf nsp12-F/7L8 (**Figure 2B**), which we suspect is due to a lower stoichiometry of the 7L8 fusion protein (**Figure 1A**, lanes 3 and 4). The RdRp complex in which nsp7 and nsp8 were co-expressed individually with nsp12 (Sf nsp12-HF/7/8) did not show measurable activity (**Supplementary Figure S2A**), probably because nsp7 and nsp8 were sub-stoichiometric (**Figure 1A**, lane 5). Pre-incubating individually expressed Sf nsp12-HF and Sf 7H8 fusion protein (1:3 ratio, **Figure 2C**) resulted in slightly less activity than the co-purified Sf nsp12-F/7H8 (**Figure 2A**). Pre-incubating the three individually expressed proteins from *E. coli* (Ec) nsp12, nsp7, and nsp8 at a ratio of 1:3:6 produced the highest activity (**Figure 2D**). Mixing Ec nsp12 with insect cell expressed 7H8 at a ratio of 1:3 (Sf 7H8, **Figure 2E**) and mixing Sf nsp12-HF with Ec nsp7 and Ec nsp8 at a ratio of 1:3:6 (**Figure 2F**) also produced moderate activity. All together, these results suggest that either co-purification of nsp12 with a nsp7-nsp8 fusion protein or mixing and preincubation of individually purified proteins, regardless of insect cell or bacterial expression, can generate active RdRp complexes.

**Figure 2.**
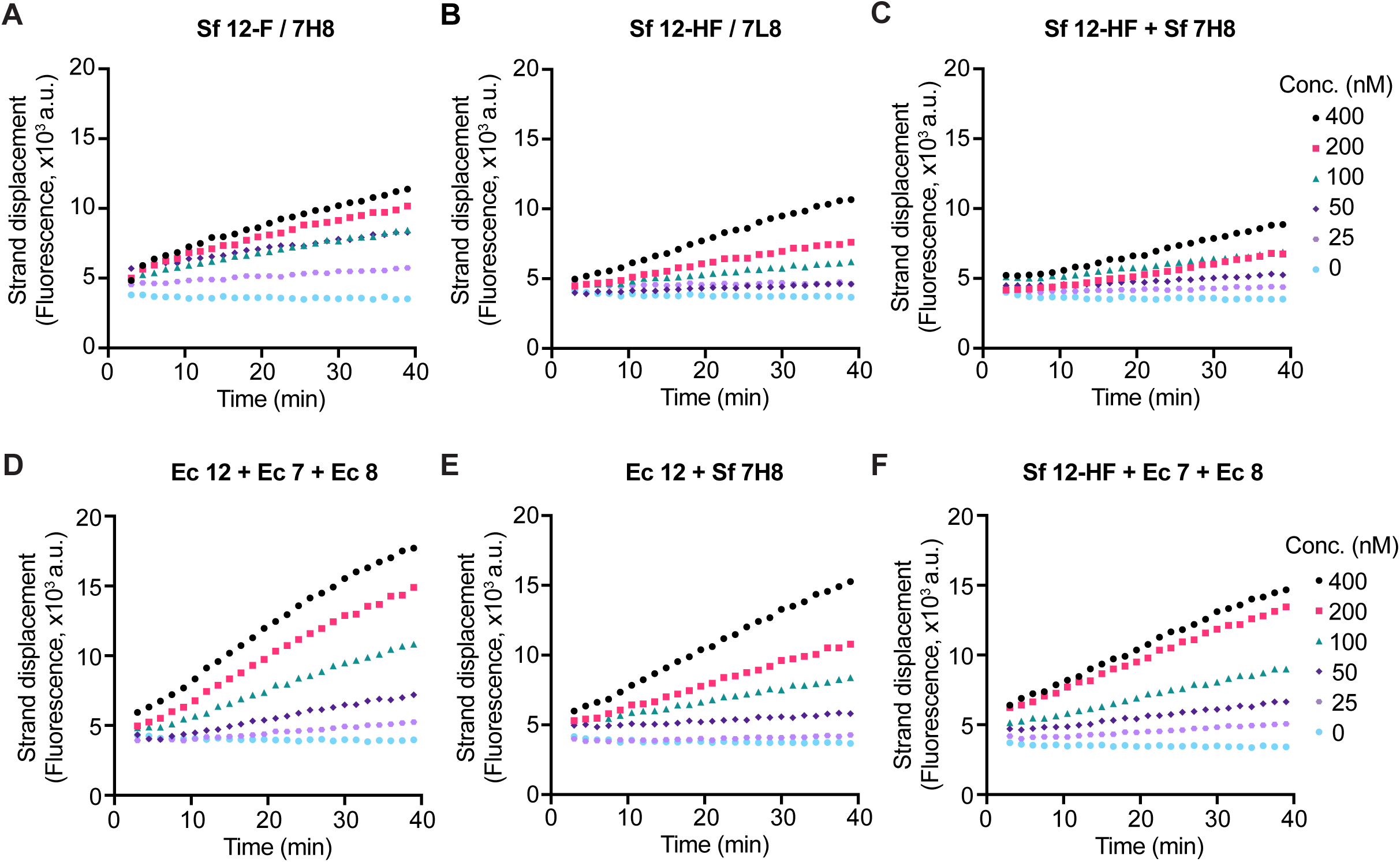
Activity of different RdRp preparations in the strand displacement assay. RdRp was mixed at the indicated concentrations with 100 nM RNA substrate and 300 µM of each NTP and Cy3 fluorescence was recorded. Enzyme preincubation was performed where indicated at 22 °C for 30 min at the indicated ratios (5 µM nsp12). (**A**) Sf nsp12-F/7H8. (**B**) Sf nsp12-HF/7L8. (**C**) Sf nsp12-HF after preincubation with Sf 7H8 (1:3 ratio). (**D**) Ec nsp12 after preincubation with Ec nsp7 and Ec nsp8 (1:3:6 ratio). (**E**) Ec nsp12 after preincubation with Sf 7H8 (1:3 ratio). (**F**) Sf nsp12-HF after preincubation with Ec nsp7 and Ec nsp8 (1:3:6 ratio).

Best protein yields were obtained in insect cells when nsp12 was co-expressed with 7L8 fusion protein (**Supplementary Table S1**), and therefore, we optimised the HTS assay conditions using this RdRp preparation (Sf nsp12-HF/7L8). The 7L8 fusion protein was sub-stoichiometric (**Figure 1A**, lane 4), so we tested if the activity of Sf nsp12-HF/7L8 could be increased by further addition of nsp12 cofactors: Ec nsp7 and Ec nsp8 or Sf 7H8. We preincubated the proteins at high concentration (10 µM for Sf nsp12-HF/7L8 either with Ec nsp7 and Ec nsp8 in 1:3:6 ratio, or with Sf 7H8 in 1:6 ratio) for 30 minutes and found they increased the activity of Sf nsp12-HF/7L8 (**Figure 3A-C**), with Sf 7H8 fusion protein addition showing the highest activity (**Figure 3C**). We also mixed Sf nsp12-HF/7L8 and Sf 7H8 at a ratio of 1:3 or 1:5 at different concentrations and found both ratios conferred similar activity at 100 nM and 150 nM Sf nsp12-HF/7L8 (**Figure 3D-E**). A final concentration of 150 nM RdRp complex in a 1:3 ratio was chosen and used for the high-throughput screen and all subsequent experiments. Finally, the presence of 5% DMSO, which is contributed by the chemical library in the high throughput screening assay, did not affect RdRp activity (**Supplementary Figure S3A**).

**Figure 3.**
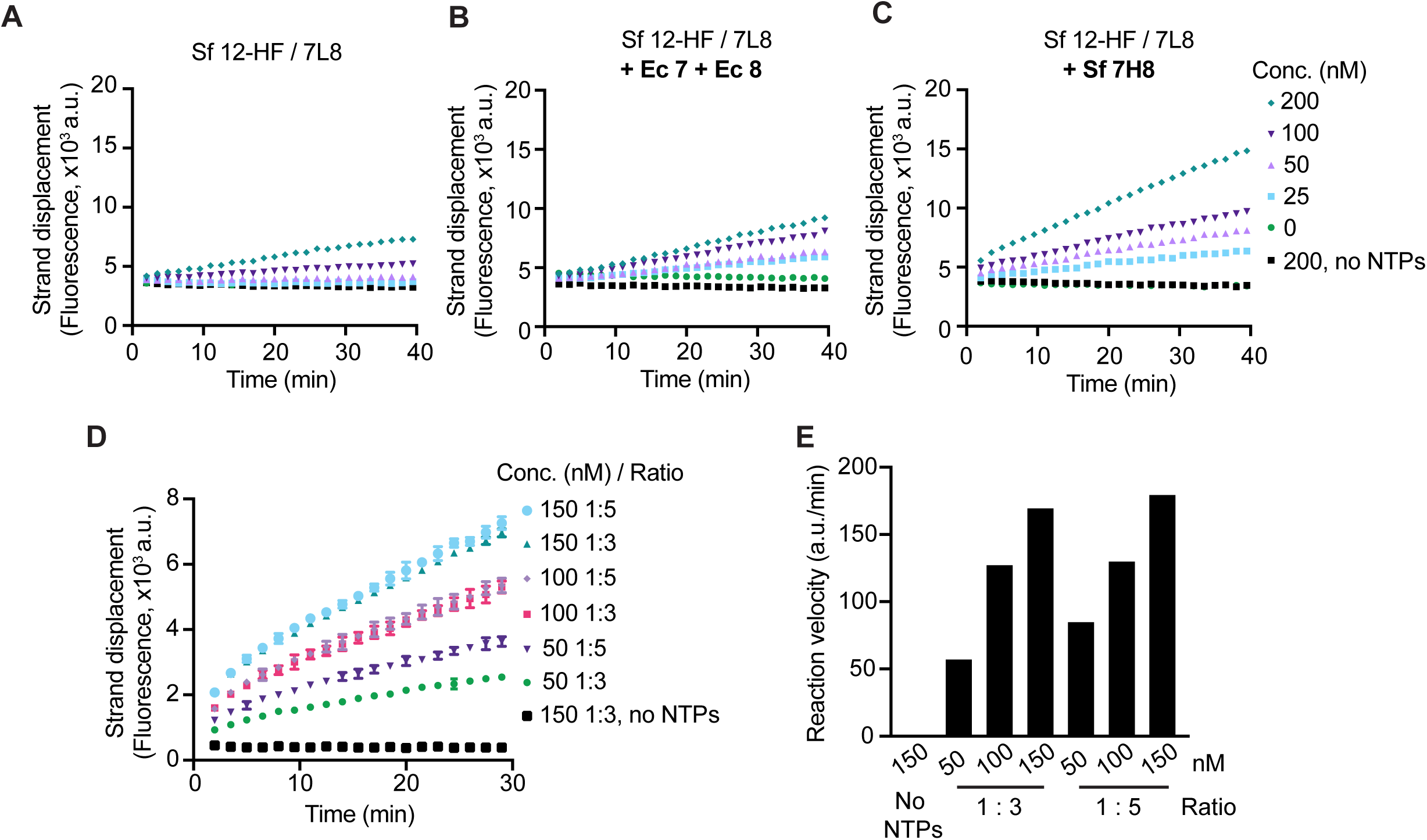
Optimisation of assay conditions for HTS. (**A-C**) Strand displacement assay using insect cell-expressed (Sf) nsp12-HF/7L8 at the indicated concentrations alone (**A**) or after preincubating at 22 °C for 30 min with Ec nsp7 and Ec nsp8 (ratio 1:3:6) (**B**) or with Sf 7H8 (ratio 1:6) (**C**). As control a condition without NTPs using 200 nM RdRp was included. (**D**) Optimization experiment to decide on RdRp concentration using HTS conditions. Sf nsp12-HF/7L8 was preincubated at 22 °C for 30 min with Sf 7H8 at either 1:3 or 1:5 ratio as specified and tested in the strand displacement assay at the indicated concentrations. (**E**) Reaction velocities extracted from the curves shown in (D).

#### Chemical library screen design and results

The optimised strand displacement assay was used to screen a custom library of over 5000 compounds for inhibitors of SARS-CoV-2 RdRp activity in a 384-well format. The compounds were incubated with RdRp at room temperature for 10 minutes, and then reactions were started by substrate addition (**Figure 4A**). Fluorescent signals were recorded in 90 second intervals for 10 cycles, which covered the linear phase of the reactions (**Supplementary Figure 4A**), and fluorescence increase over time was used to calculate the reaction velocity for each well. The library was screened at two compound concentrations, 1.25 µM (**Figure 4B**) and 3.75 µM (**Figure 4C**). Reactions that showed >15% reduction in the velocity (**Figure 4B,C**) or >10% reduction in the endpoint signal (**Supplementary Figure S4C,D**) were manually inspected (example in **Figure 4D**). 64 compounds were selected as primary hits. Compounds were then discounted if: i) the compound also scored as a strong hit in parallel HTSs of other SARS-CoV-2 enzymes (with the exception of sulphated naphthylamine derivatives, see below) (Biochem J, this issue); ii) the compound was reported or strongly predicted to be a promiscuous inhibitor due to colloidal aggregation (LogP value >3.6 and >85% similarity to a known aggregator) (45, 46); iii) the compound was reported or predicted to be a nucleic acid intercalator. This analysis resulted in the selection of 18 compounds for further validation experiments (**Supplementary Table S2**). Notably, 5 compounds of the 18 selected primary hits could be categorised as sulphated naphthylamine derivatives (suramin, NF 023, PPNDS, Evans Blue and Diphenyl Blue). These compounds were also identified as inhibitors of SARS-CoV-2 nsp13 helicase in a parallel screen (Zeng et al., Biochem J, this issue). Suramin and suramin-like molecules are polyanionic compounds that likely bind to positively charged patches found in a diverse array of proteins (47–53), including nucleic acid binding proteins like helicases (54) and viral polymerases (55–59). *In vitro* validation experiments described in Zeng et al. showed that all five compounds inhibit SARS-CoV-2 nsp13 helicase with IC_50_ values between 0.5 - 6 µM and SARS-CoV-2 RdRp with IC_50_ values between 0.5 – 5 µM. They are not predicted to have aggregation tendencies (**Supplementary Table S2**) and their nsp13 inhibition was shown to be insensitive to detergent (Zeng et al, Biochem J, this issue), making nonspecific inhibition due to colloidal aggregation unlikely (60). Suramin, as the main representative drug of this group, was selected to be included in further *in vitro* validation experiments as part of this study. Ultimately, 14 compounds were chosen for further validation experiments.

**Figure 4.**
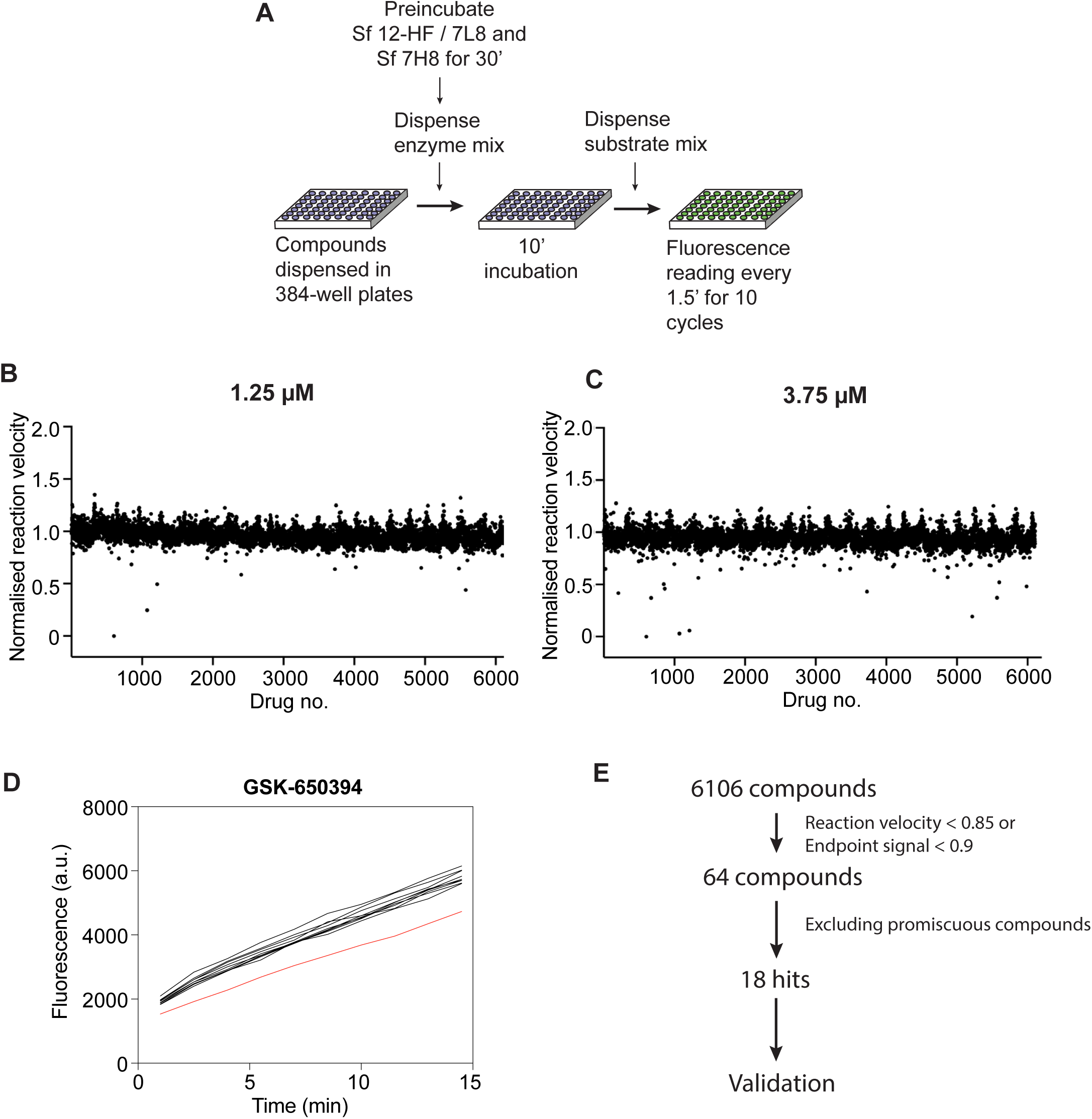
High-throughput chemical library screen against SARS-CoV-2 RNA-dependent RNA polymerase. (**A**) Logistics of the screen. A custom chemical library consisting of over 5000 compounds was screened against RdRp activity using the FRET-based strand displacement assay in a 384-well format. RdRp was prepared by preincubation of Sf nsp12-HF/7L8 and Sf 7H8 in a 1:3 ratio for 30 min at room temperature. RdRp was dispensed into compound-containing 384-well plates and incubated for 10 min. Reactions were started by addition of a substrate mix and florescence monitored in 90 s intervals. (**B-C**) Results of the HTS screen performed at 1.25 µM (B) and 3.75 µM (C) compound concentration. The normalised reaction velocity plotted against compound number is shown. (**D**) Kinetic curves with >15% reduction in reaction velocity or >10% reduction in fluorescent signal at the endpoint were inspected manually. As example, kinetic data for compound GSK-650394 is shown (red curve, data from surrounding wells in black). (**E**) Summary of the HTS hit selection strategy. From over 5000 compounds tested in the screen, 64 were considered primary hits after manual inspection of HTS reactions, which showed a reduction of reaction velocity below 85%, or a reduction of endpoint signal below 90%. Out of these, 46 primary hits were eliminated as they likely represent nonspecific modes of enzymatic inhibition such as colloidal aggregation or interference with the substrate structure. As part of this analysis promiscuous compounds that were identified as hits in other SARS-CoV-2 HTS (Biochem J, this issue) were removed with the exception of 5 suramin and suramin-like compounds, which were also identified in the SARS-CoV-2 nsp13 helicase HTS (Zeng et al., Biochem J, this issue). *In vitro* validation of the effect of suramin and suramin-like compounds on the activity of SARS-CoV-2 helicase and SARS-CoV-2 RdRp can be found in Zeng et al. (this issue). A total of 18 compounds (including 5 suramin and suramin-like compounds) were selected as hits, of which 14 were included in further *in vitro* validation in this work.

### Validation of hits

In order to confirm and quantify RdRp inhibition by the selected compounds *in vitro*, we performed compound titration experiments to determine the drug concentration at which half maximal inhibition is achieved (IC_50_). We carried these experiments out in the presence and in the absence of the non-ionic detergent Triton X-100 to rule out unspecific inhibition due to compound aggregation (60). We found that 6 compounds showed only weak or no clear RdRp inhibition (**Supplementary Figure S5A, Supplementary Table S3**). Another 2 compounds showed interference with the fluorescent substrate by quenching (**Supplementary Figure S5A, Supplementary Table S3**). Therefore, these compounds were excluded from further analysis. The remaining 6 compounds inhibited RdRp with IC_50_ values of 0.43 µM - 56 µM and showed no or only minor sensitivity to detergent, indicating they might be *bona fide* RdRp inhibitors (**Figure 5A, Table 1**).

**Figure 5.**
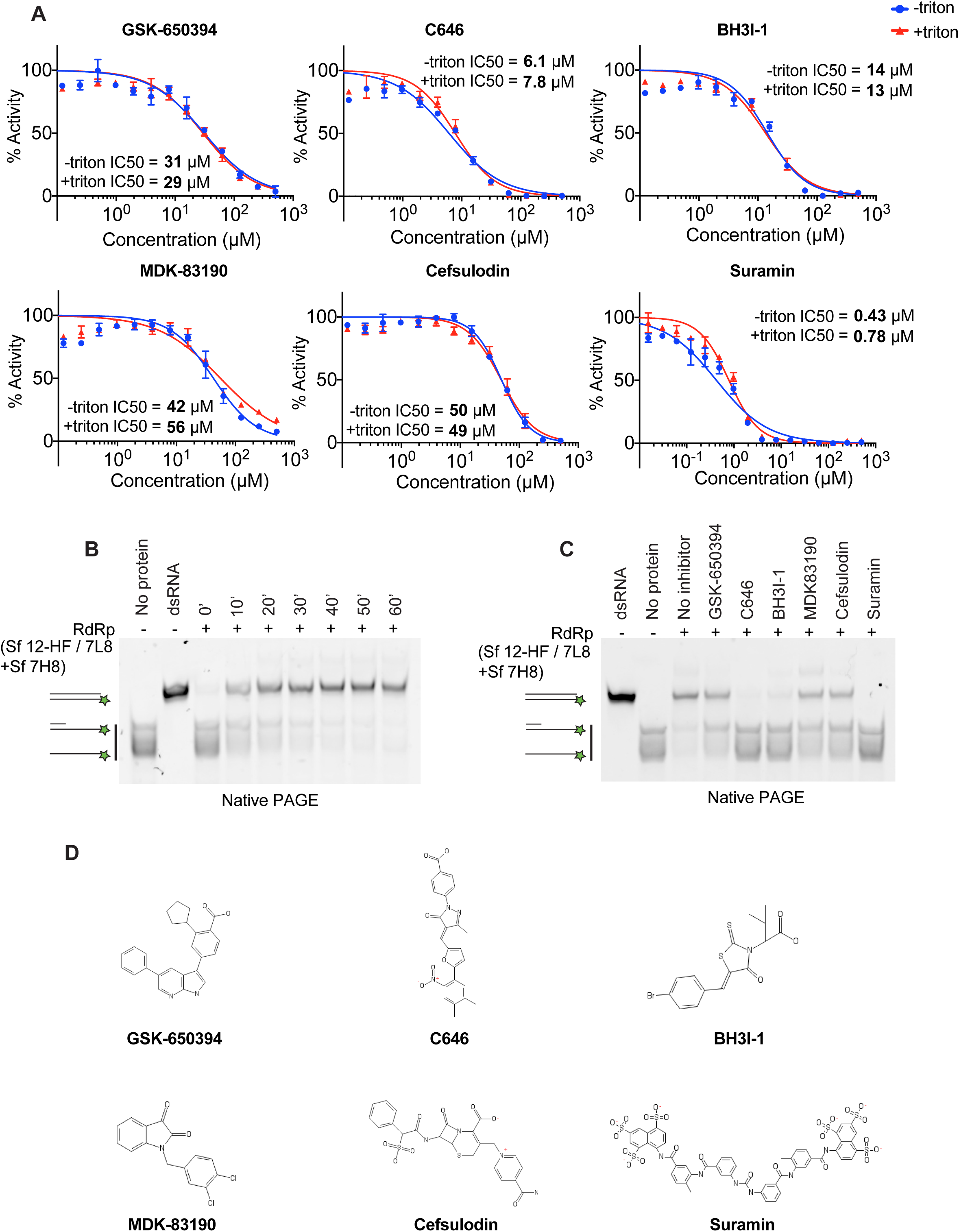
*In vitro* validation of selected compounds identified as RdRp inhibitors. (**A**) Concentration-response curves of selected compounds using the strand displacement assay. The experiment was performed using 150 nM RdRp, 100 nM RNA substrate and 300 µM of each NTP in the presence (+Triton) or absence (-Triton) of 0.01% Triton X-100. Quenching controls are shown in **Supplementary Figure S5B**. IC_50_ values were calculated using Prism software. (**B-C**) Native gel-based assays using a primed RNA substrate as in Figure 1C. Reactions were started by mixing 300 nM RdRp complex (formed by preincubation of Sf nsp12-HF/7L8 and Sf 7H8 at 1:3 ratio) with 50 nM RNA substrate and 1 mM NTPs. Reaction products were analysed by native PAGE and visualisation of Cy3 fluorescence. (**B**) RdRp reactions were incubated for the indicated amounts of time. (**C**) Validation of selected compounds using 30-minute RdRp reactions. RdRp complex was incubated with 50 µM of the indicated compounds for 10 minutes before reactions were started by substrate addition. Controls: a preformed dsRNA with the same size as the reaction product (dsRNA), the primed substrate (No protein). (**D**) Chemical structures of selected RdRp inhibitors.

**Table 1.** Inhibitory activity of compounds against SARS-CoV-2 RdRp in vitro.

| Compound name | IC<sub>50</sub> (μM) |  | Data figure |
| --- | --- | --- | --- |
|  | No detergent | + 0.01% Triton X-100 |  |
| Suramin<sup>a</sup> | 0.43 | 0.78 | 4 |
| C646 | 6.1 | 7.8 | 4 |
| BH3I-1 | 14 | 13 | 4 |
| GSK-650394 | 31 | 29 | 4 |
| MDK-83190 | 42 | 56 | 4 |
| Cefsulodin | 50 | 49 | 4 |

<sup>a</sup> For a further characterisation of suramin and the suramin-related compounds NF 023, PPNDS, Evans Blue, Diphenyl Blue see Zeng et al., Biochem J (this issue), 2021.

We tested the 6 compounds in the gel-based polymerisation assay which monitors RNA synthesis more directly using the same primed RNA substrate as in **Figure 1C**. The RdRp preparation used here (Sf 12-HF/7L8 + Sf 7H8, ratio 1:3) also showed time-dependent RNA duplex formation (**Figure 5B**). We performed a 30-minute reaction with the compounds at 50 µM and found they inhibited RdRp consistent with their IC_50_ values from the kinetic assay. Suramin, C646 and BH3I-1 inhibited RNA duplex formation almost completely, and the compounds with higher IC_50_ values, GSK-650394, MDK83190 and Cefsulodin showed partial inhibition (**Figure 5C**).

### Viral inhibition assays

The validation experiments indicated that 6 compounds are genuine SARS-CoV-2 RdRp inhibitors in biochemical assays (**Figure 5A** and **Table 1**). We therefore evaluated their potential antiviral activity against SARS-CoV-2 in Vero E6 cells. Cells were treated with the compounds and then infected with SARS-CoV-2. After 22 hours, viral plaques were analysed by viral nucleocapsid (N) protein immunofluorescence (**Figure 6A**). We tested the 5 novel RdRp inhibitors for their potential antiviral activity using this assay (**Figure 6B**). Dose–response curves and the half maximal effective concentration (EC_50_) for each compound was calculated (**Figure 6C**). Two of them, GSK-650394 and C646, presented a dose-dependent reduction in the levels of SARS-CoV-2 N protein (green) in Vero E6 cells with low micromolar EC_50_ values (EC_50_=7.6 µM and EC_50_=19 µM respectively, **Figure 6B,C**). Suramin antiviral activity in Vero E6 cells following the same aforementioned protocol has been evaluated by us in the accompanying manuscript Zeng et al. and an EC_50_ of 9.9 µM against SARS-CoV-2 was observed (Zeng et al, Biochem J, this issue).

**Figure 6.**
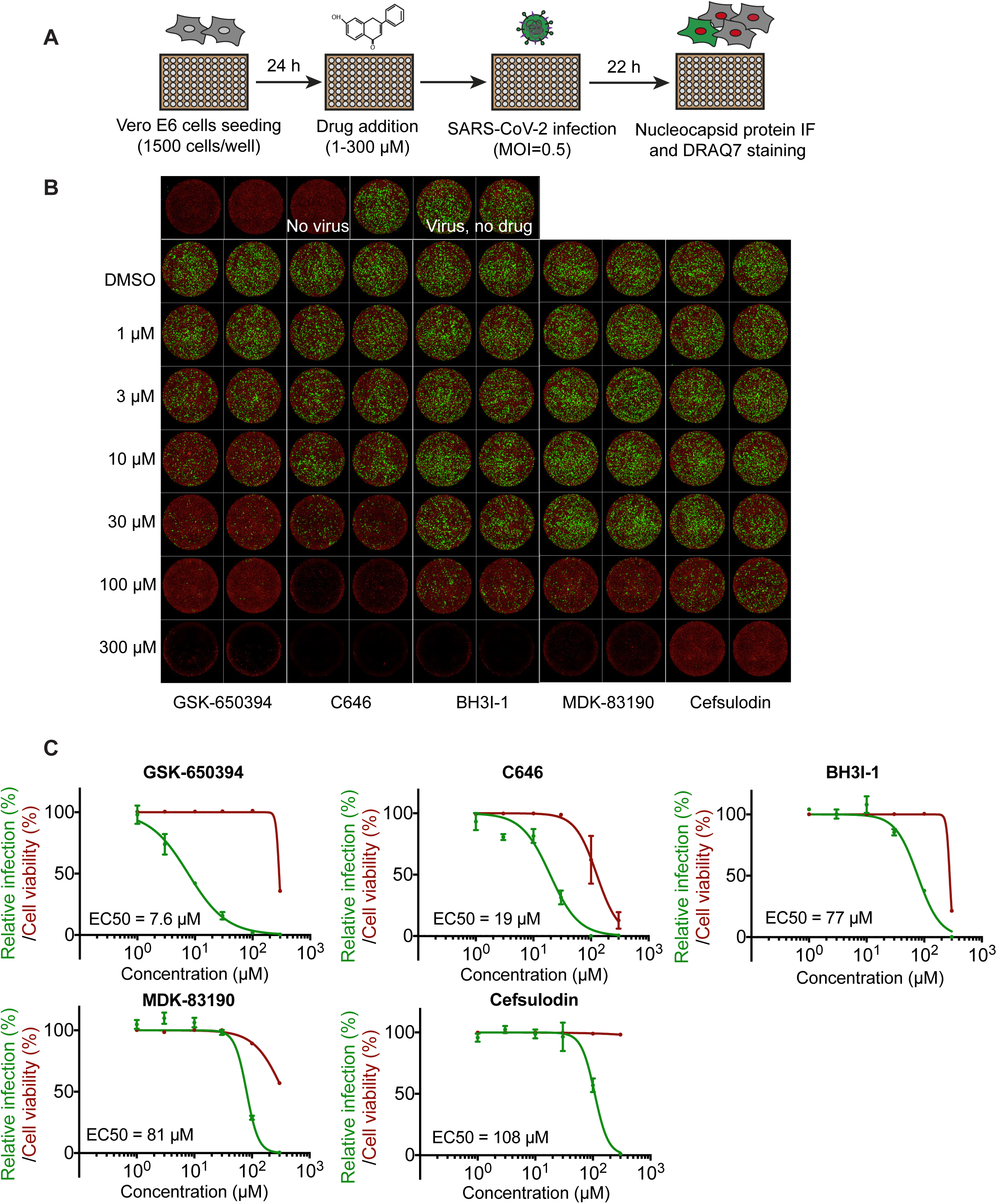
Antiviral activity of selected compounds against SARS-CoV-2 in Vero E6 cells. (**A**) Viral inhibition experiments workflow graphical representation. In brief, 24 hours after Vero E6 cells were seeded in a 96-well format, they were treated with selected drugs at specified concentrations. Then, cells were infected with a SARS-CoV-2 isolate at a MOI of 0.5 PFU per cell. Twenty-two hours later, cells were fixed and analysed by immunofluorescence staining and imaging. (**B**) SARS-CoV-2 antiviral activity of GSK-650394, C646, BH3I-1, MDK-83190 and Cefsulodin. Representative overlaid images of viral N protein immunofluorescence (green) and DNA dye DRAQ7 staining (red). (**C**) Dose-response curve analysis. Viral infection values represent the area of viral plaques visualised by viral nucleocapsid protein staining (green curves) and cell viability was measured in the same experiment as the area of cells stained with the DNA dye DRAQ7 (red curves). Data is plotted as percentage relative to DMSO only control wells (100%). Data represent the mean and standard deviation (SD) of 3 replicates. Areas were calculated using FIJI and EC_50_ values were calculated using Prism software.

Combination therapy, the use of two or more antivirals with different modes of action, is a known strategy for limiting drug resistance as well as achieving better physiological outcomes (61). The latter is usually caused by reduced toxicity due to the use of lower doses in combination protocols compared to monotherapy doses (61). Consequently, a potential synergy between RdRp inhibitors with different mechanisms of action (non-nucleoside analogues identified here and a nucleoside analog, remdesivir) was assessed (38, 62, 63). Following the viral inhibition protocol described in **Figure 6A**, we performed dose-response experiments for each compound in the presence of a fixed concentration of 0.5 µM remdesivir, which as a single agent inhibited viral infection by less than 10%. Under these assay conditions, none of the validated inhibitors showed significant synergy with remdesivir (**Supplementary Figure S6**).

## Discussion

The replication/transcription complex (RTC) of coronaviruses catalyses the synthesis of all genomic RNAs (viral replication) and sub-genomic RNAs (viral transcription). The core of SARS-CoV-2 RTC is the RNA-dependent RNA polymerase holoenzyme, composed of nsp12, the catalytic subunit, together with its cofactors nsp7 and nsp8 (19–21, 64). SARS-CoV-2 RdRp is the target of remdesivir, the only antiviral approved for human use to treat COVID-19 by regulatory agencies (62). Remdesivir is a broad-spectrum nucleoside analog first used for treatment of Ebola infections (4). Remdesivir’s moderate success in COVID-19 treatment is still a matter of controversy as new clinical trials suggest that it is not as effective as first thought (7–10). Coronaviruses harbour a unique RNA-replication proofreading trait conferred by the 3’-to-5’ exonuclease activity of nsp14 (29), offering a possible explanation as to why the development or repurposing of effective nucleoside analogues, with the exception of remdesivir, has been unsuccessful (31, 36). Therefore, the characterisation of non-nucleoside analog-type drugs that target SARS-CoV-2 RdRp might be needed more than ever. Moreover, COVID-19 is the third known transfer, after SARS-CoV in 2003 and MERS-CoV in 2012, of an animal coronavirus to a human population in only 20 years and therefore it is highly probable that new zoonotic outbreaks will occur in the future (65). Thus, it is vital to develop numerous broad-spectrum antiviral strategies, that together with vaccination programs, could curb future cross-species transmissions.

Here, we have expressed, purified and characterised a diverse array of SARS-CoV-2 RdRp complexes. We used two expression hosts, a bacterial system (*E. coli*) and a baculovirus-insect cell system and showed that recombinant proteins from both systems individually expressed or expressed as a complex lead to active proteins. We characterised different constructs of nsp12, nsp7 and nsp8 and tested activities of RdRp complexes formed with different stoichiometries of the subunits. We showed that the SARS-CoV-2 RdRp can strand displace downstream RNA encountered during synthesis. Based on this characteristic of SARS-CoV-2 RdRp, we designed a novel and robust FRET-based assay that allows real-time monitoring of RdRp activity *in vitro*. We utilised this assay to screen a custom chemical library to identify compounds that inhibit SARS-CoV-2 RdRp enzymatic activity. Our study identified GSK-650394, C646 and BH3I-1 as well as confirmed suramin and suramin-like compounds as *in vitro* inhibitors of SARS-CoV-2 RdRp activity. These compounds do not have typical chemical structures of nucleoside analogues and thus likely inhibit RdRp through a different mechanism of action from remdesivir. Further work is required to understand their mode of inhibition. Cryo-electron microscopy structures of SARS-CoV-2 RdRp have been presented by several groups (20, 38, 40, 64, 66, 67). Future compound-protein structures may help reveal the inhibitory mechanisms of the RdRp inhibitors identified in this study.

GSK-650394, suramin and C646 showed also significant inhibition on viral growth in cell-based assays with an EC_50_ of 7.6 µM, 9.9 µM and 19 µM respectively (**Figure 6B,C**) (nsp13 paper, Biochem J, this issue). **GSK-650394** is an inhibitor of SGK-1 kinase (68) that plays an important role in a diverse range of cellular processes such as stress response, ion transport, inflammation and cell proliferation (69). GSK-650394 has been reported to impede growth of cellular models of cancers (68, 70) and has also been implicated in the inhibition of replication of influenza A virus, an enveloped negative-sense ssRNA virus, in human A549 cells (71). In our study GSK-650394 showed a low micromolar EC_50_ of 7.6 µM in inhibiting SARS-CoV-2 in cell-based assays, with effects on cell death only at the highest concentration tested (300 µM). **Suramin** is an antiparasitic drug first introduced by Bayer in 1916 that is still used to treat sleeping sickness and river blindness (72). Suramin has been repurposed for a number of diverse applications including the inhibition of purinergic signaling, treatment of viral infections and treatment for advanced malignancies reflecting its multitude of targets (47). Suramin is a polyanionic compound and probably binds tightly to positively charged patches found in DNA or RNA binding proteins (47, 48). This is consistent with our findings, as suramin and suramin analogues inhibit *in vitro* both SARS-CoV-2 RdRp and SARS-CoV-2 nsp13 helicase (Zeng et al, Biochem J, this issue). **C646** is a competitive inhibitor of the histone acetyltransferase p300 that amongst other effects results in a reduction of pro-inflammatory gene expression in cells and animal models (73–76). C646 was also shown to suppress the replication of different strains of influenza A viruses in A549 cells with an EC_50_ of 16 μM by affecting several steps of the viral life cycle (77). Murine models of influenza infection treated with C646 also showed significant reduction in viral replication with no associated toxicity (77). Further *in vitro*, cell-based and pre-clinical studies are needed to confirm and characterise these novel RdRp inhibitors. Moreover, they can be used as promising lead compounds to design improved drug candidates.

In summary, we have identified small-molecule *non-*nucleoside analog inhibitors of SARS-CoV-2 RdRp that also inhibited viral replication in cell-based assays. We provide evidence that GSK-650394, suramin and C646 could be considered as drug candidates that deserve further evaluation, as these compounds were found to exhibit antiviral activity against SARS-CoV-2 in a relevant cell culture model at non-cytotoxic concentrations. This screening strategy can be expanded to other small molecule compound libraries and together with the biochemical insights regarding SARS-CoV-2 RdRp expression and purification described here, these should accelerate the search for improved antivirals that target SARS-CoV-2 RdRp holoenzyme.

## Materials and Methods

### Protein expression

The following proteins were expressed in baculovirus-infected insect cells: nsp7-His_6_-nsp8 (Sf 7H8), nsp12-His_6_-3xFlag (Sf nsp12-HF), nsp12-3xFlag/nsp7-His_6_-nsp8 (Sf nsp12-F/7H8), nsp12-His_6_-3xFlag/nsp7-(GGS)_2_-nsp8 (Sf nsp12-HF/7L8), and nsp12-His_6_-3xFlag/nsp7/nsp8 (Sf nsp12-HF/7/8). The coding sequences of SARS-CoV-2 nsp7, nsp8 and nsp12 (NCBI reference sequence NC_045512.2) were codon-optimised for *S. frugiperda* and synthesized (GeneArt, Thermo Fisher Scientific). Baculoviral expression constructs were generated using the biGBac vector system (78). An empty polyhedrin (polh) expression cassette from pLIB was inserted into the vectors pBIG1a, pBIG1b, and pBIG1c to generate pBIG1a(polh), pBIG1b(polh), and pBIG1c(polh).

Nsp12 was subcloned into pBIG1a(polh) either to contain a C-terminal His_6_-3xFlag tag (protein sequence: M-nsp12-GGSHHHHHHGSDYKDHDGDYKDHDIDYKDDDDK, pBIG1a_nsp12-His_6_-3xFlag) or to contain a C-terminal 3xFlag tag (protein sequence: M-nsp12-GGSDYKDHDGDYKDHDIDYKDDDDK, pBIG1a_nsp12-3xFlag). Nsp7 and nsp8 were either subcloned individually into pBIG1b(polh) and pBIG1c(polh) to not contain a tag (pBIG1b_nsp7 and pBIG1c_nsp8), or into pBIG1b(polh) to be expressed as fusion protein with a His_6_-linker (protein sequence: M-nsp7-HHHHHH-nsp8, pBIG1b_nsp7-His_6_-nsp8) or with a (GGS)_2_-linker (protein sequence: M-nsp7-GGSGGS-nsp8, pBIG1b_nsp7-(GGS)_2_-nsp8). Co-expression constructs were generated by subcloning expression cassettes from pBIG1 vectors into pBIG2abc vectors: pBIG1a_nsp12-3xFlag, pBIG1b_nsp7-His_6_-nsp8 and empty pBIG1c were used to generate pBIG2abc_nsp12-3xFlag/nsp7-His_6_-nsp8 (nsp12-F/7H8). The vectors pBIG1a_nsp12-His_6_-3xFlag, pBIG1b_nsp7-(GGS)_2_-nsp8, and empty pBIG1c were used to generate pBIG2abc_nsp12-His_6_-3xFlag/nsp7-(GGS)_2_-nsp8 (nsp12-HF/7L8). The vectors pBIG1a_nsp12-His_6_-3xFlag, pBIG1b_nsp7 and pBIG1c_nsp8 were used to generate pBIG2abc_nsp12-His_6_-3xFlag/nsp7/nsp8 (nsp12-HF/7/8). Baculoviruses were generated and amplified in Sf9 insect cells (Thermo Fisher Scientific) using the EMBacY baculoviral genome (79). For protein expression, Sf9 cells were infected with baculovirus and collected 48 hours after infection, flash-frozen and stored at −70°C.

The following proteins were expressed in *Escherichia coli* (Ec): 14His-SUMO-nsp7 (Ec 7), 14His-SUMO-nsp8 (Ec 8) and 14His-SUMO-nsp12 (Ec 12). The coding sequences of SARS-CoV-2 nsp7, nsp8 and nsp12 were codon-optimised for *E. coli* and synthesized (Genewiz). Nsp7, nsp8 and nsp12 were subcloned into the K27SUMO expression vector (41) to code for Ulp1-cleavable 14His-SUMO fusion proteins. For expression the vectors were transformed into T7 Express lysY competent *E. coli* cells (New England Biolabs). Expression cultures were grown in LB containing 50 µg/ml kanamycin to an OD_600_ of 0.5 and expression was induced with 0.4 mM IPTG at 16°C. After overnight incubation, cells were collected, flash-frozen and stored at −70°C.

### Protein purification

All proteins were purified at 4°C. The proteins Sf nsp12-F/7H8 and Sf nsp12-HF/7/8 were purified by FLAG affinity, Heparin affinity and size exclusion chromatography. Insect cell pellets containing these proteins were resuspended in Flag buffer (50 mM HEPES pH 7.4, 150 mM KCl, 5 mM MgCl_2_, 10% glycerol, 1 mM DTT) supplemented with protease inhibitors (Roche Complete Ultra EDTA-free tablets, 10 µg/ml leupeptin, 10 µg/ml pepstatin A, 1 mM AEBSF) and lysed by Dounce homogenization. After centrifugation at 39000g, 60 min, 4°C, the protein was purified from the cleared lysate by affinity to Anti-FLAG M2 Affinity gel (Sigma-Aldrich) and eluted in Flag buffer containing 0.5 mg/ml 3xFLAG peptide. The Flag eluate was diluted 1:6 with dilution buffer (50 mM HEPES pH 7.4, 5 mM MgCl_2_, 10% glycerol, 1 mM DTT) and loaded onto a HiTrap Heparin HP column (GE Healthcare) equilibrated in Heparin buffer A (50 mM HEPES pH 7.4, 25 mM KCl, 5 mM MgCl_2_, 10% glycerol, 1 mM DTT). The protein was eluted with a gradient to Heparin buffer B (Heparin buffer A containing 500 mM KCl). Protein-containing fractions were concentrated and further purified by gel filtration on a Superdex 200 Increase 10/300 GL column (GE Healthcare) in Heparin buffer A containing 150 mM KCl. The protein was concentrated, flash-frozen and stored at −70°C.

The proteins Sf 7H8, Sf nsp12-HF, and Sf nsp12-HF/7L8 were purified by Ni-NTA affinity, ion exchange, and size exclusion chromatography. Insect cell pellets containing these proteins were resuspended in His-tag buffer (50 mM Tris-HCl pH 7.5, 500 mM NaCl, 5 mM MgCl_2_, 10% glycerol, 1 mM DTT, 0.02% Nonidet P40 substitute, 20 mM imidazole) containing protease inhibitors (Roche Complete Ultra EDTA-free tablets, 10 µg/ml leupeptin, 10 µg/ml pepstatin A, 1 mM AEBSF) and lysed using a Dounce homogenizer. The lysate was cleared by centrifugation at 39000g, 60 min, 4°C and the supernatant added to Ni-NTA Agarose (Invitrogen) equilibrated in His-tag buffer. The beads were washed with His-tag buffer and protein was eluted with His-elution buffer (50 mM Tris-HCl pH 7.5, 200 mM NaCl, 5 mM MgCl_2_, 10% glycerol, 1 mM DTT, 0.02% Nonidet P40 substitute, 200 mM imidazole). The eluate was diluted 1:5 with MonoQ buffer A (20 mM Tris-HCl pH 8.0, 2 mM MgCl_2_, 10% glycerol, 0.02% Nonidet P40 substitute, 1 mM DTT) and loaded onto a MonoQ 5/50 GL column (GE Healthcare) equilibrated with MonoQ buffer A containing 50 mM NaCl. After gradient elution to 1 M NaCl, protein-containing fractions were concentrated and further purified by gel filtration using a Superdex 200 Increase 10/300 GL column (GE Healthcare). 7H8 was purified in SEC-150 buffer (25 mM HEPES pH 7.5, 150 mM NaCl, 2 mM MgCl_2_, 10% glycerol, 0.02% Nonidet P40 substitute, 1 mM DTT). The proteins nsp12-HF and nsp12-HF/7L8 were purified in SEC-300 buffer (25 mM HEPES pH 7.5, 300 mM NaCl, 2 mM MgCl_2_, 10% glycerol, 0.02% Nonidet P40 substitute, 1 mM DTT). The proteins were concentrated, flash-frozen and stored at −70°C.

The proteins 14His-SUMO-nsp7, 14His-SUMO-nsp8, and 14His-SUMO-nsp12 were purified by Ni-NTA affinity, proteolysis by His-Ulp1, removal of cleaved 14His-SUMO and His-Ulp1 by Ni-NTA, ion exchange and size exclusion chromatography. *E. coli* pellets containing these proteins were resuspended in His-SUMO buffer (50 mM Tris-HCl pH 7.5, 500 mM NaCl, 5 mM MgCl_2_, 10% glycerol, 1 mM DTT, 0.05% Nonidet P40 substitute, 30 mM imidazole) containing protease inhibitors (Roche Complete EDTA-free tablets, 10 µg/ml leupeptin, 10 µg/ml pepstatin A, 1 mM AEBSF) and were lysed by sonication. The lysate was cleared by centrifugation at 39000g, 60 min, 4°C and the protein was purified from the supernatant using Ni-NTA Agarose (Invitrogen). The beads were washed with His-SUMO buffer and protein was eluted with His-SUMO elution buffer (50 mM Tris-HCl pH 7.5, 300 mM NaCl, 5 mM MgCl_2_, 10% glycerol, 1 mM DTT, 0.05% Nonidet P40 substitute, 400 mM imidazole). The SUMO-specific protease His-Ulp1 was added to the eluates to a concentration of 0.02 mg/ml and the proteins were dialysed overnight using His-SUMO dialysis buffer (25 mM Tris-HCl pH 7.5, 250 mM NaCl, 5 mM MgCl_2_, 10% glycerol, 0.02% Nonidet P40 substitute, 1 mM DTT). Cleaved 14His-SUMO and His-Ulp1 were removed by affinity to Ni-NTA Agarose (Invitrogen). The flowthrough was diluted 1:5 with MonoQ buffer A and loaded onto a MonoQ 5/50 GL column (GE Healthcare), which was equilibrated with MonoQ buffer A containing 50 mM NaCl. After gradient elution to 1 M NaCl, all fractions were analysed, and protein-containing fractions concentrated. The proteins were further purified by gel filtration using a Superdex 200 Increase 10/300 GL column (GE Healthcare). nsp7 and nsp8 were purified in SEC-150 buffer, and nsp12 was purified in SEC-300 buffer. The proteins were concentrated, flash-frozen and stored at −70°C.

### FRET-based strand displacement assay

A FRET-based fluorescence-quenching assay was designed to monitor strand displacement activity of the RdRp complex. The assay uses a tripartite RNA substrate. HPLC-purified RNA oligonucleotides were purchased from Eurofins Genomics or Integrated DNA Technologies with the following sequences:

Cy3 strand: 5’-Cy3-GCACUUAGAUAUGACUCGUUCUGCAGGCCAGUUAA-3’

Quencher strand: 5’-CUGCGUCUAAGUGC-(AbRQ)-3’

Primer strand: 5’-UUAACUGGCC-3’

The Cy3 strand, the quencher strand and the primer strand were annealed at 1:2:2 ratio at a concentration of 10 µM:20 µM:20 µM respectively by heating the oligonucleotide mix to 75°C for 5 minutes and gradually cooling it down to 4°C over 50 minutes in annealing buffer (10 mM Tris-HCl pH 8.0, 1 mM EDTA, 20 mM NaCl and 5 mM MgCl_2_). A reaction was typically started at room temperature by adding 10 µL of a 2x substrate solution containing the RNA substrate and NTPs in reaction buffer (20 mM HEPES pH 7.5, 10 mM KCl, 1 mM DTT, 5 mM MgCl_2_, 2 mM MnCl_2_, 0.1 mg/ml BSA, 0.01% Triton X-100) to 10 µL of a 2x enzyme solution containing the RdRp complex in reaction buffer. A typical reaction contained a final concentration of 100 nM RNA substrate, 300 µM of each NTP, and 150 nM RdRp. Upon displacement of the quencher strand by the RdRp complex during synthesis, Cy3 is no longer quenched and thus able to fluoresce. The fluorescent signal was read at 545 nm (excitation) and 575 nm (emission) with 10 nm bandwidth using a Tecan Spark microplate reader.

### FRET-based polymerization assay

The Cy3 template strand and the primer were annealed at 1:2 ratio using the protocol described above. A reaction was typically started at room temperature by adding 10 µL of a 2x substrate solution containing the RNA substrate and NTPs in reaction buffer to 10 µL of a 2x enzyme solution containing the RdRp complex in reaction buffer. The quencher strand was added at 2-fold concentration (relative to the Cy3 strand) after a 60-min reaction. The quencher strand was allowed to anneal with any unpolymerized template RNA for 35 min at room temperature before the fluorescent signal was read.

### Gel-based polymerization assay

The Cy3 template strand and the primer were annealed at 1:2 ratio using the protocol described above. A reaction was typically started at room temperature by adding 5 µL of a 2x substrate solution containing the RNA substrate and NTPs in reaction buffer to 5 µL of a 2x enzyme solution containing the RdRp complex in reaction buffer. The reaction was stopped by adding 5 µL stopping buffer (20 mM EDTA, 0.5% SDS, 2 mg/ml proteinase K, 10% glycerol) and incubating at 37°C for 5 min. Reaction products were then analysed by running on a 20% non-denaturing PAGE gel in 0.5X TBE buffer (Life Technologies) for 20 min at room temperature and 200 V. An Amersham Imager 600 was used to image Cy3 fluorescence. To test compound inhibition, a compound volume of 0.5 µL at a concentration of 1 mM was incubated with 5 µL 2x enzyme solution for 10 min before starting the reaction.

### RdRp high-throughput screening assay

The screening plates containing the custom library of compounds were prepared and stored as described in (Zeng et al, Biochem J, this issue). For the HTS assay, first, 10 µM of Sf nsp12-HF/7L8 complex was incubated with 30 µM Sf 7H8 for 30 min on ice to form the RdRp complex. Then the RdRp complex was diluted in reaction buffer, before being dispensed at 10 µL volume to the plates to incubate with compounds for 10 min at room temperature. To start the reaction, 10 µL 2x substrate mix (600 µM NTPs, 200 nM RNA substrate) in reaction buffer was dispensed and the plates were spun briefly before being transferred to a Tecan Spark microplate reader. The fluorescent signal was first read at 1 min of reaction and then read every 1.5 min for 10 cycles. The screening was performed twice, one with a final compound concentration of 1.25 µM and one with 3.75 µM.

### Data analysis

MATLAB was used to process data. The slope of fluorescent signal increase over time was calculated using a linear regression model for each compound as the reaction velocity (V). Then V for each compound is normalised against the DMSO controls in each plate as following: Normalised_V=V/mean(V_DMSO), Where mean(V_DMSO) is the average of velocity values of DMSO controls in a particular plate. The endpoint signal of each compound was also normalised against DMSO controls. Both the normalised reaction velocity and endpoint signal were used for hit selection. Compounds that gave >15% reduction in the reaction velocity or >10% reduction in the endpoint signal were further inspected. A kinetic curve (example in Figure 4D), in which the y-axis shows fluorescent signals, and the x-axis shows timepoints, was plotted for each of the compounds that passed the threshold. 10 wells that were before and after the compound well were plotted together in the same graph. We manually inspected the curves and confirmed that 64 compounds gave obvious inhibition.

The Z-score was also calculated for each compound (Supplementary Figure S4B) as following: Z-score=[Normalised_V -mean(Normalised_V_all)]/SD(Normalised_V_all), Where mean(Normalised_V_all) is the average of normalised velocity values of all compounds and SD(Normalised_V_all) is the standard deviation of normalised velocity values of all compounds.

To assess the quality of the screen, the Z-factor was calculated for each plate as following: Z-factor=1 – [3*SD(V_DMSO) + 3*SD(background)]/[mean(V_DMSO) – mean(background) Where SD(V_DMSO) is the standard deviation of reaction velocity values of all DMSO controls in a particular plate, SD(background) is the standard deviation of reaction velocity values of background wells containing no enzyme, and mean(background) is the average of reaction velocity values of background wells. The average Z-factor for the screen is 0.71, indicating the screen is good, as described by (80).

The method used for predicting drug aggregation propensity is described in (46) and can be accessed in http://advisor.bkslab.org/.

### Tested Drugs

Compounds selected for *in vitro* experimental validation and cell-based studies were purchased and resuspended following manufacturer’s instructions. See **Supplementary Table S4** for more details.

### Viral inhibition assay

The production of the SARS-CoV-2 virus isolate used in these assays and the in-house generation of the SARS-CoV-2 nucleocapsid (N) recombinant antibody is described elsewhere (Biochem J, nsp13 paper). The conditions for the viral inhibition assay were the following: 15.000 Vero E6 cells (NIBC, UK) resuspended in DMEM containing 10% FBS were seeded into each well of 96-well imaging plates (Greiner 655090) and cultured overnight at 37°C and 5% CO2. The next day, 5x solutions of compounds were generated by dispensing 10 mM stocks of compounds into a V-bottom 96-well plate (Thermo 249946) and back filling with DMSO to equalise the DMSO concentration in all wells using an Echo 550 (Labcyte) before resuspending in DMEM containing 10% FBS. The assay plates with seeded Vero E6 cells had the media replaced with 60 µl of fresh growth media, then 20 µl of the 5x compounds were stamped into the wells of the assay plates using a Biomek Fx automated liquid handler. Finally, the cells were infected by adding 20 µl of SARS-CoV-2 with a final MOI of 0.5 PFU/cell. 22 h post infection, cells were fixed, permeabilised, and stained for SARS-CoV-2 N protein using Alexa488-labelled-CR3009 antibody and cellular DNA using DRAQ7 (Abcam). Whole-well imaging at 5x was carried out using an Opera Phenix (Perkin Elmer) and fluorescent areas and intensity calculated using the Phenix-associated software Harmony (Perkin Elmer).

## Data Availability Statement

All data associated with this paper will be deposited in FigShare (https://figshare.com/).

## Author Contributions

**Agustina P. Bertolin**: Conceptualization, Methodology, Validation, Formal analysis, Investigation, Writing - Original Draft, Writing - Review & Editing, Visualization. **Florian Weissmann**: Conceptualization, Methodology, Validation, Formal analysis, Investigation, Writing - Original Draft, Writing - Review & Editing, Visualization. **Jingkun Zeng**: Conceptualization, Methodology, Software, Validation, Formal analysis, Investigation, Data Curation, Writing - Original Draft, Writing - Review & Editing, Visualization. **Viktor Posse**: Conceptualization, Methodology, Validation, Investigation, Writing - Review & Editing. **Jennifer Milligan:** Investigation, Resources. **Berta Canal**: Conceptualization, Investigation, Writing - Review & Editing. **Rachel Ulferts**: Investigation. **Mary Wu**: Investigation. **Lucy S. Drury**: Resources. **Michael Howell**: Resources, Supervision. **Rupert Beale**: Resources, Supervision. **John FX Diffley:** Conceptualization, Methodology, Formal analysis, Writing - Review & Editing, Supervision, Project administration, Funding acquisition.

## Acknowledgements

We thank Anne Early for assistance. This work was supported by the Francis Crick Institute, which receives its core funding from Cancer Research UK (FC001066), the UK Medical Research Council (FC001066), and the Wellcome Trust (FC001066). This work was also funded by a Wellcome Trust Senior Investigator Award (106252/Z/14/Z) to J.F.X.D. FW and BC have received funding from the European Union’s Horizon 2020 research and innovation programme under the Marie Skłodowska-Curie grant agreement Nos 844211 and 895786. JZ has received funding from a Ph.D. fellowship awarded by Boehringer Ingelheim Fonds.

## Supplementary Figure legends

**Supplementary Figure S1.**
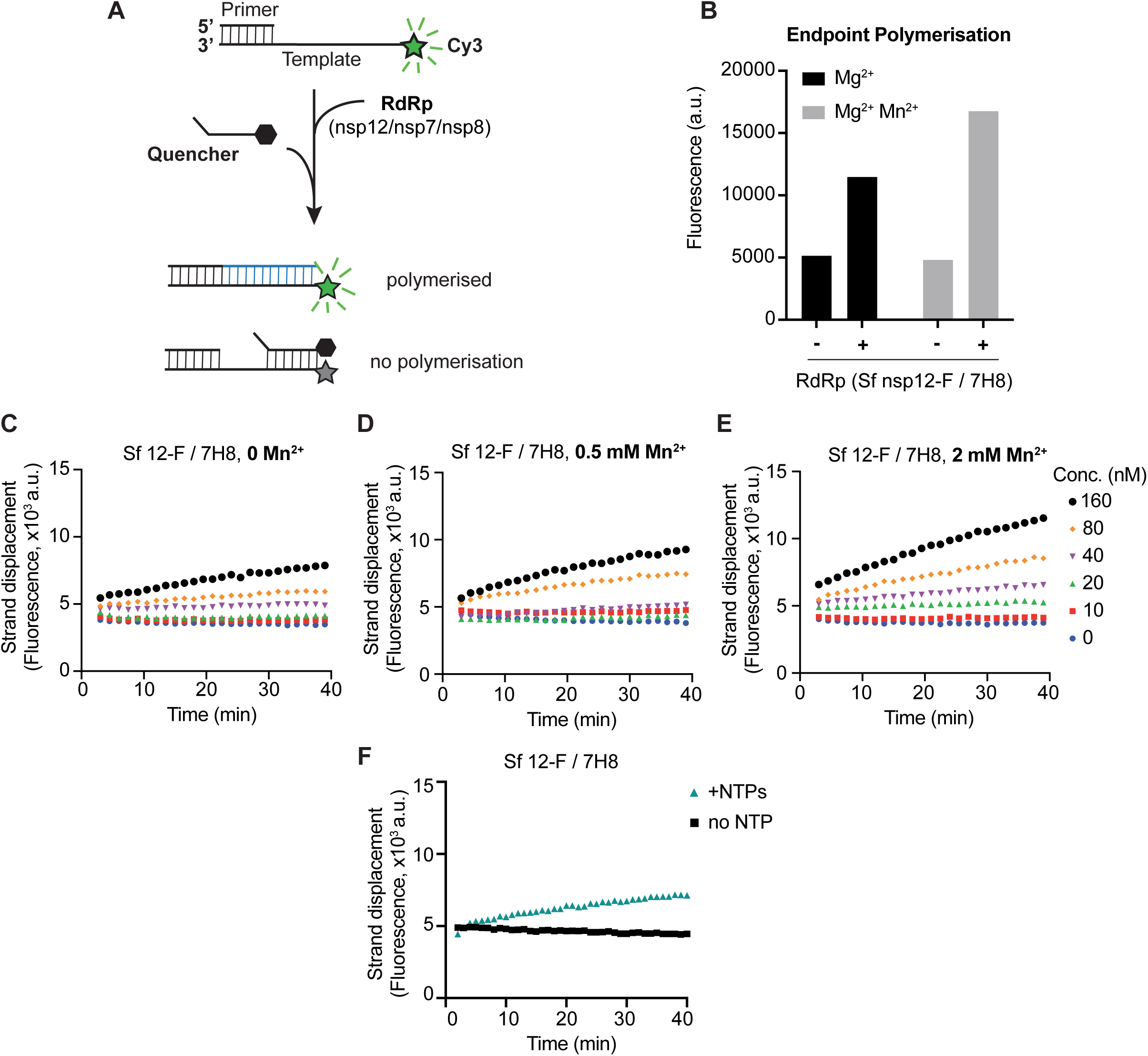
(**A**) Schematic diagram illustrating the FRET-based primer extension assay. The RNA substrate is composed of a Cy3 fluorophore-containing template strand and an annealed primer strand. The substrate is incubated with RdRp, which extends the primer strand generating duplex RNA. After incubation, a quencher strand is added, which can only anneal to the template strand and quench Cy3 fluorescence if RNA synthesis did *not* occur. (**B**) Results of the FRET-based primer-extension assay shown in (A). The quencher strand was added after 60 min of RdRp and primed-substrate incubation and fluorescence was measured after another 35 min. (**C-E**) Strand displacement assay using the indicated concentrations Sf nsp12-F/7H8 in the absence (C) or presence of 0.5 mM Mn^2+^ (D) or 2 mM Mn^2+^ (E). (**F**) Strand displacement assay using Sf nsp12-F/7H8 in the presence (+NTPs) or absence (no NTP) of 300 µM NTPs.

**Supplementary Figure S2.**
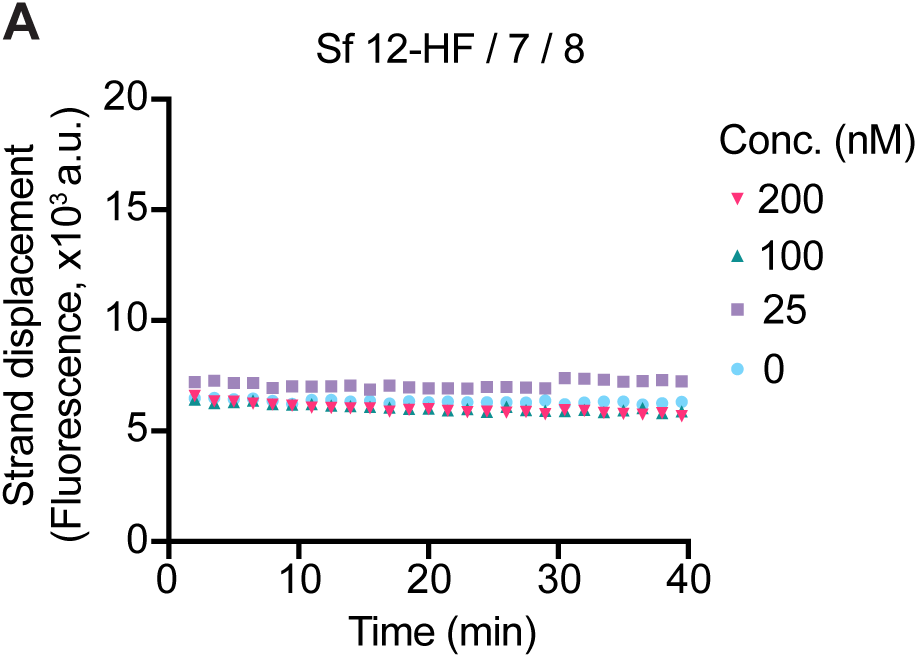
(**A**) Strand displacement assay using the indicated concentrations of insect cell (Sf)-expressed nsp12-HF/7/8. This experiment was performed in parallel with the experiment shown in Figure 1E.

**Supplementary Figure S3.**
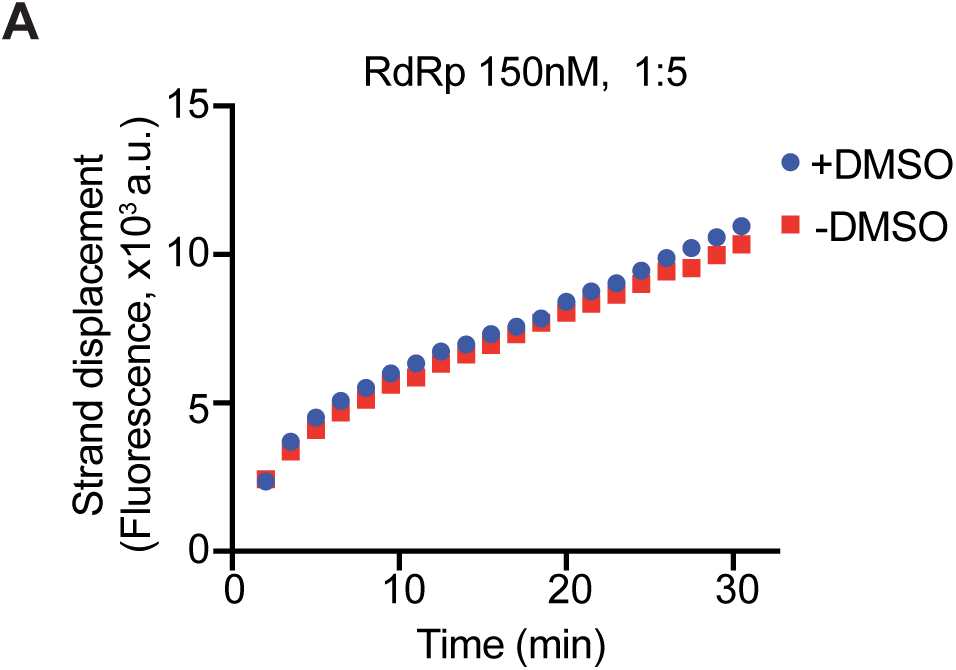
(**A**) Strand displacement assay using 150 nM Sf nsp12-HF/7L8 premixed with Sf 7H8 (1:5 ratio) in the presence (+) or absence (−) of 5% DMSO.

**Supplementary Figure S4.**
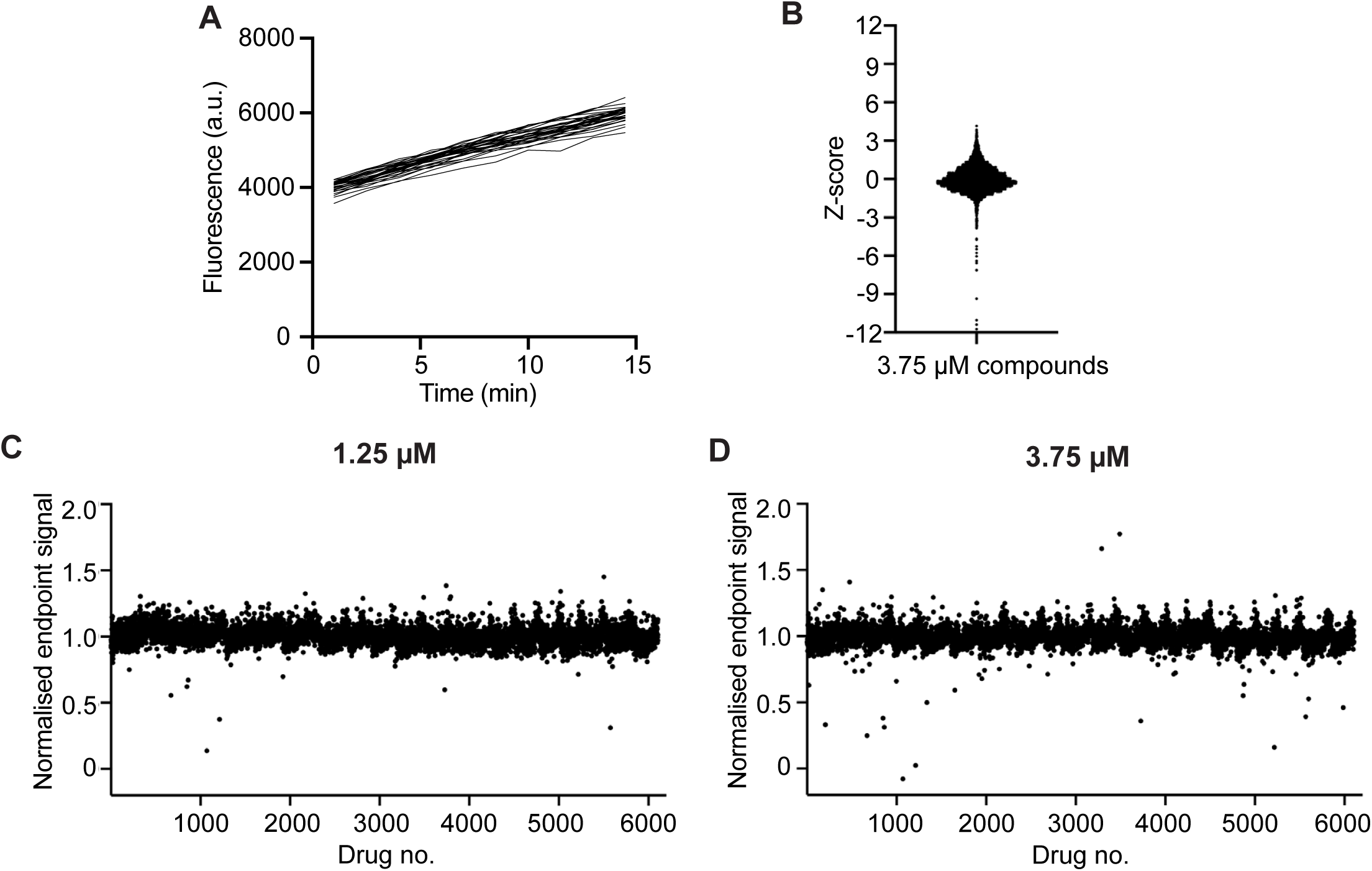
High-throughput SARS-CoV-2 RdRp inhibitor screen. (**A**) Representative kinetic data from control reactions in the HTS screen. For screen analysis, reaction velocities were extracted from the slope. (**B**) Z-score distribution for the screen performed at 3.75 µM compound concentration. (**C-D**) Analysis of the screen by endpoint signal. Normalised endpoint signal is plotted against compound number for the screen performed at 1.25 µM (C) or 3.75 µM (D) compound concentration.

**Supplementary Figure S5.**
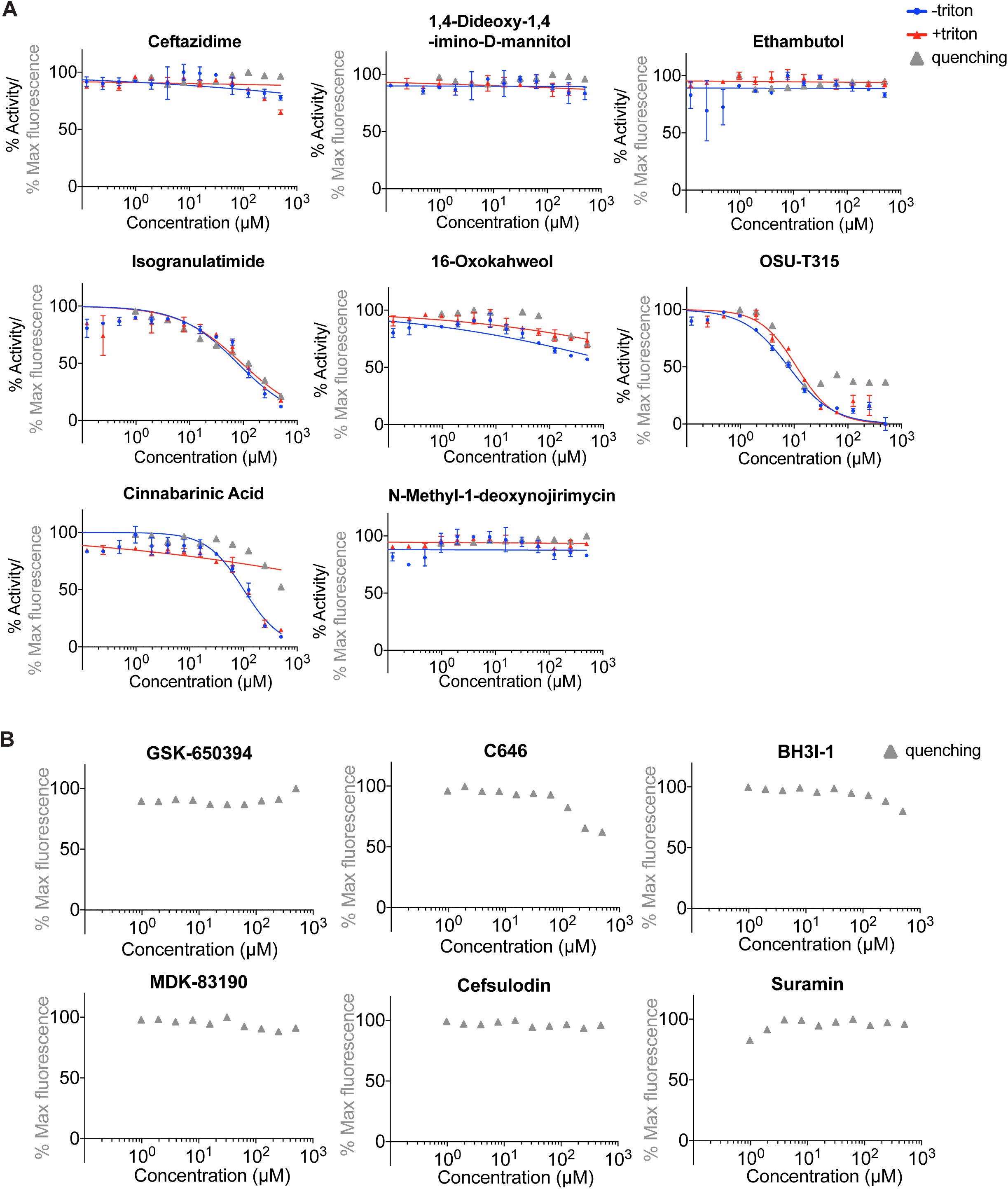
Concentration-response curves for validation of selected compounds identified in SARS-CoV-2 RdRp screen. (**A**) The experiment was performed using 150 nM RdRp, 100 nM RNA substrate, 300 µM of each NTP and in the presence (+Triton) or absence (- Triton) of 0.01% Triton X-100. Fluorescence quenching properties of the compounds were tested to identify false positives that interfered with the fluorescent substrate (quenching). (**B**) Quenching controls for the experiment shown in Figure 5A. IC_50_ values were calculated using Prism software.

**Supplementary Figure S6.**
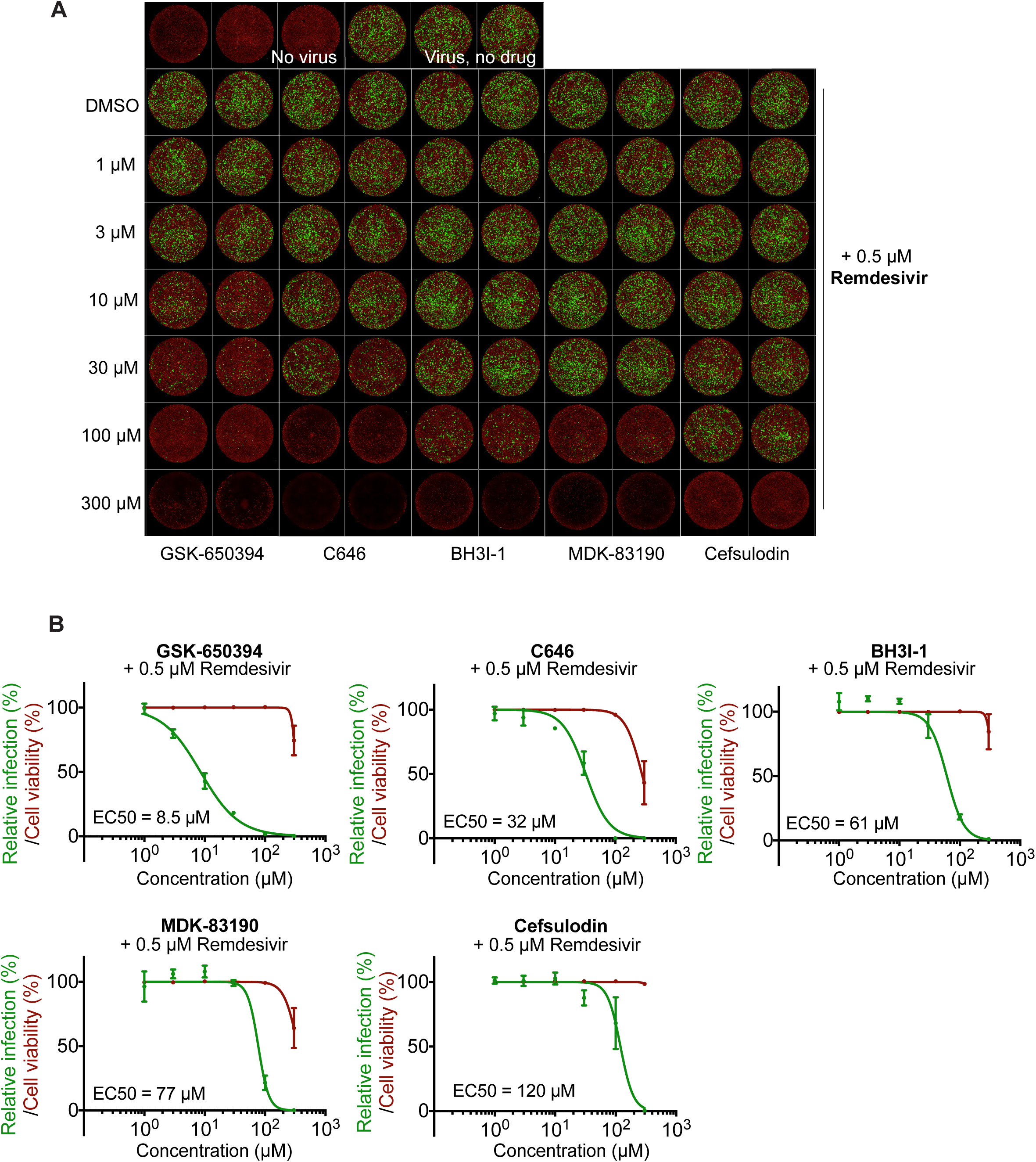
Comparative dose–response curves of selected antiviral compounds against SARS-CoV-2 in cell culture. **(A)** Anti-SARS-CoV-2 activities of GSK-650394, C646, BH3I-1, MDK-83190 and Cefsulodin in combination with 0.5 µM remdesivir following protocol detailed in **Figure 6A**. Representative overlaid images showing N protein immunofluorescence (green) and DRAQ7 nuclei staining (red). **(B)** Dose-response curve analysis. Viral infection values represent the area of virus infected cells visualised by N protein staining (green curves) and, simultaneously, cell viability was measured as the area of cells stained for DRAQ7 (red curves). Data is plotted as percentage normalised to 0.5 µM remdesivir only treated wells (100%). Values represent mean and standard deviation (SD) of 3 replicates. Areas were calculated using FIJI software and EC_50_ values were calculated using Prism software.

**Supplementary Table S1.** Overview of purified SARS-CoV-2 RdRp proteins including yields.

| Protein preparation | Expression system | Yield (mg) | Yield (nmol) | Construct | Purification |
| --- | --- | --- | --- | --- | --- |
| **7H8** | Baculovirus-insect cell | 19.5<sup>a</sup> | **607<sup>a</sup>** | nsp7-His6-nsp8 | Ni-NTA, MonoQ, S200 |
| **nsp12-HF** | Baculovirus-insect cell | 1.92 | 17.3 | nsp12-His6-3xFlag | Ni-NTA, MonoQ, S200 |
| **nsp12-F/7H8** | Baculovirus-insect cell | 0.084 | 0.59 | nsp12-3xFlag/nsp7-His6-nsp8 | FLAG M2, Heparin, S200 |
| **nsp12-HF/7L8** | Baculovirus-insect cell | 17.0<sup>a</sup> | **119<sup>a</sup>** | nsp12-His6-3xFlag/nsp7-GGSGGS-nsp8 | Ni-NTA, MonoQ, S200 |
| **nsp12-HF/7/8** | Baculovirus-insect cell | 0.053 | 0.37 | nsp12-His6-3xFlag/nsp7/nsp8 | FLAG M2, Heparin, S200 |
| **nsp7** | *E. coli* | 1.96 | 213 | 14His-Sumo-nsp7 | Ni-NTA, Ulp1, Ni-NTA, MonoQ, S200 |
| **nsp8** | *E. coli* | 5.34 | 244 | 14His-Sumo-nsp8 | Ni-NTA, Ulp1, Ni-NTA, MonoQ, S200 |
| **nsp12** | *E. coli* | 0.65 | 6.1 | 14His-Sumo-nsp12 | Ni-NTA, Ulp1, Ni-NTA, MonoQ, S200 |

<sup>a</sup> Combined yield from 2 preparations.

**Supplementary Table S2.** Predicted aggregation propensity of 18 HTS hit compounds.

| Compound name | LogP | Structural Similarity Index |
| --- | --- | --- |
| NF 023 | -6.1 | - |
| Suramin | -5.7 | - |
| Cefsulodin | -5.7 | - |
| Ceftazidime | -5 | - |
| PPNDS | -3.3 | - |
| Evans Blue | -2.7 | - |
| Diphenyl Blue | -2.7 | - |
| 1,4-Dideoxy-1,4-imino-D-mannitol | -2.6 | - |
| N-Methyl-1-deoxynojirimycin | -2.2 | - |
| Ethambutol | 0.4 | - |
| Cinnabarinic Acid | 1.1 | - |
| Isogranulatimide | 2 | - |
| BH3I-1 | 3.1 | - |
| 16-Oxokahweol | 3.3 | - |
| MDK 83190 | 3.8 | - |
| C646 | 4.7 | - |
| OSU-T315 | 5.2 | - |
| GSK-650394 | 5.8 | - |

Compounds analysed using the open access tool Aggregator Advisor See (Irwin et al., 2015)

**Supplementary Table S3.** Compounds that showed only weak or no clear activity against SARS-CoV-2 RdRp in vitro or interfered with the fluorescent substrate by quenching.

| Compound name | IC<sub>50</sub> (μM) |  | Data figure |
| --- | --- | --- | --- |
|  | No detergent | + 0.01% Triton X-100 |  |
| Cinnabarinic acid | >100 | >100 | S4A |
| Ceftazidim | no clear inhibition | no clear inhibition | S4A |
| 1,4-Dideoxy-1,4-imino-D-mannitol | no clear inhibition | no clear inhibition | S4A |
| Ethambutol | no clear inhibition | no clear inhibition | S4A |
| 16-Oxokahweol | no clear inhibition | no clear inhibition | S4A |
| N-Methyl-1-deoxynojirimycin | no clear inhibition | no clear inhibition | S4A |
| OSU-T315 | <sup>a</sup> | <sup>a</sup> | S4A |
| Isogranulatimide | <sup>a</sup> | <sup>a</sup> | S4A |

<sup>a</sup> IC<sub>50</sub> could not be assessed due to compound-related quenching effects on fluorescent substrate

**Supplementary Table S4.** Purchased drugs for *in vitro* validation and viral inhibition experiments.

| Compound name | CAS Number | Company | Catalog Number |
| --- | --- | --- | --- |
| Suramin | 129-46-4 | Sigma | S2671 |
| Cefsulodin | 52152-93-9 | Sigma | C 8145 |
| Cinnabarinic Acid | 606-59-7 | Sigma | SML0096 |
| 1,4-Dideoxy-1,4-imino-D-mannitol | 114976-76-0 | Sigma | D 8390 |
| C646 | 328968-36-1 | Sigma | SML0002 |
| OSU-T315 | 2070015-22-2 | MedChem Express | HY-18676 |
| MDK 83190 | 79183-19-0 | MedChem Express | HY-18633 |
| Ceftazidime | 72558-82-8 | MedChem Express | HY-B0593 |
| BH3I-1 | 300817-68-9 | MedChem Express | HY-100383 |
| Remdesivir | 1809249-37-3 | MedChem Express | HY-104077 |
| Isogranulatimide | 244148-46-7 | Calbiochem/Millipore | 371957 |
| 16-Oxokahweol | 108664-99-9 | Cayman Chemicals | CAY30117 |
| N-Methyl-1-deoxynojirimycin | 69567-10-8 | Biovitica | BVT-0130-M001 |
| Ethambutol | 1070-11-7 | Selleck Chemicals | S4004 |
| GSK-650394 | 890842-28-1 | Tocris | 3572 |

